# A deep dilated convolutional residual network for predicting interchain contacts of protein homodimers

**DOI:** 10.1101/2021.09.19.460941

**Authors:** Raj S. Roy, Farhan Quadir, Elham Soltanikazemi, Jianlin Cheng

## Abstract

**Motivation:** Deep learning has revolutionized protein tertiary structure prediction recently. The cutting-edge deep learning methods such as AlphaFold can predict high-accuracy tertiary structures for most individual protein chains. However, the accuracy of predicting quaternary structures of protein complexes consisting of multiple chains is still relatively low due to lack of advanced deep learning methods in the field. Because interchain residue-residue contacts can be used as distance restraints to guide quaternary structure modeling, here we develop a deep dilated convolutional residual network method (DRCon) to predict interchain residue-residue contacts in homodimers from residue-residue co-evolutionary signals derived from multiple sequence alignments of monomers, intrachain residue-residue contacts of monomers extracted from true/predicted tertiary structures or predicted by deep learning, and other sequence and structural features.

**Results:** Tested on three homodimer test datasets (Homo_std dataset, DeepHomo dataset, and CASP14-CAPRI dataset), the precision of DRCon for top L/5 interchain contact predictions (L: length of monomer in a homodimer) is 43.46%, 47.15%, and 24.81% respectively, which is substantially better than two existing deep learning interchain contact prediction methods. Moreover, our experiments demonstrate that using predicted tertiary structure or intrachain contacts of monomers in the unbound state as input, DRCon still performs reasonably well, even though its accuracy is lower than when true tertiary structures in the bound state are used as input. Finally, our case study shows that good interchain contact predictions can be used to build high-accuracy quaternary structure models of homodimers.

**Availability:** The source code of DRCon is available at https://github.com/jianlin-cheng/DRCon.

## 1 Introduction

Proteins fold into three-dimensional (3D) structures to carry out biological functions such as catalyzing chemical reactions and transporting nutrients. The 3D structure of a single protein chain is called tertiary structure. The tertiary structures of multiple protein chains usually interact to form a complex structure (i.e., quaternary structure). Both tertiary structure and quaternary structure are important for protein function. Because the experimental determination of protein structure is low-throughput and can be applied to only a small portion of proteins in the nature, the computational prediction of protein tertiary and quaternary structure is critical for obtaining structural information for most proteins to study their function.

The computational methods for predicting protein tertiary structures and quaternary structures are periodically evaluated in the Critical Assessment of Protein Structure Prediction (CASP) (Kryshtafovych et al., 2014; Moult et al., 2016; Kryshtafovych et al., 2019; Kwon et al., 2021) and the Critical Assessment of Protein Interaction (CAPRI) (Lensink et al., 2016, 2018, 2021), respectively, or the joint experiment of the two. Driven by the application of deep learning methods to predicting residue-residue contacts and distances (Wang et al., 2017; Adhikari et al., 2018; Jones & Kandathil, 2018; Li et al., 2019; Hou et al., 2020; Senior et al., 2020; Yang et al., 2020; Wu et al., 2021) in the last several years, tertiary structure prediction has reached unprecedented high accuracy. In the 2020 CASP14 experiment, AlphaFold2 (Jumper et al., 2021) predicted high-quality structures for most CASP14 targets with the accuracy equal to or close to that of the experimental structure determination. Recently, AlphaFold2 was applied to predict the structures for all the proteins in several species including human (Tunyasuvunakool et al., 2021).

Despite the drastic advance in protein tertiary structure prediction, the prediction of quaternary structure has progressed slowly and still cannot reach high accuracy for most protein complexes. One reason is more effort has been put into tertiary structure prediction than quaternary structure prediction because the former is needed as input for the latter. Another reason is the application of deep learning methods to protein quaternary structure prediction is still in the early stage and much fewer deep learning methods for quaternary structure prediction than tertiary structure prediction have been developed.

The most common approach to quaternary structure prediction is classic protein docking algorithms (Gray et al., 2003; Lyskov & Gray, 2008; Venkatraman et al., 2009; Li & Kihara, 2012; Pierce et al., 2014; Johansson-Åkhe et al., 2020), leveraging the geometric and electrostatic complementarity between protein tertiary structures. The residue-residue coevolutionary methods such as the direct coupling analysis (DCA) (Hopf et al., 2015; Ovchinnikov et al., 2014) that were originally designed to predict intrachain residue-residue contacts in a protein chain were also used to predict interchain contacts from multiple sequence alignments (MSAs) of protein complex (e.g., protein heterodimers). The DCA based methods require a large number of sequences in MSAs to generate accurate interchain contact predictions, which are not available for most protein complexes because there are not many known protein complexes available. The problem is alleviated for protein homodimers (a protein complex consisting of two identical chains) because the MSA of a monomer (a single chain) in a homodimer contains both intrachain and interchain residue-residue co-evolutionary signals (Quadir et al., 2021). The advantage of using the MSA of a monomer is that it is generally much deeper than the MSA of a protein complex. Recently several deep learning methods such as DNCON2_Inter (Quadir et al., 2021) and DeepHomo (Yan & Huang, 2021) use the MSA of a monomer in a homodimer to predict interchain contacts in homodimers.

Another interesting recent development is the application of AlphaFold2 and RoseTTAFold (Baek et al., 2021) - the two cutting-edge deep learning methods designed for prediction of tertiary structure to the prediction of the quaternary structures of several protein complexes, demonstrating the great potentials of deep learning methods for predicting protein quaternary structures. However, because the two methods are not specially designed for quaternary structure prediction and are not trained on the protein complex data, there is a significant need to develop more deep learning methods directly targeting quaternary structure prediction.

In this work, we develop a dilated convolutional residual neural network called DRCon to directly predict interchain contacts in homodimers from the MSA, intrachain contacts, and other features of the monomers forming the homodimers. We test our method rigorously on the CASP14-CAPRI dataset, DeepHomo test dataset and also on Homo_std test dataset. It performs better than two other deep learning methods (DeepHomo and DNCON2_Inter) for interchain contact prediction. The method works not only with true tertiary structures of monomers in the bound state as input but also predicted tertiary structures of monomers in the unbound state (e.g., tertiary structure models predicted by AlphaFold2). Moreover, we demonstrate that good interchain contact predictions can be used to build high-quality quaternary structures of homodimers.

## 2 Materials and Methods

### 2.1 Datasets

Two residues from the two chains in a homodimer are considered an interchain contact if the Euclidean distance between any two heavy atoms of the two residues is less than or equal to 6Å (Ovchinnikov et al., 2014; Quadir et al., 2021; Zhou et al., 2018). Multiple homodimer datasets with known quaternary structures and interchain contacts are used to develop DRCon. The Homo_std dataset used in DNCON2_Inter is used to train, validate, and test DRCon. Homo_std was derived from the homodimers in the 3D Complex database (Levy et al., 2006). All the complexes of the 3D Complex were released before October of 2005. The dimers in the database whose two chains have >= 95% sequence identity are treated as homodimers to create Homo_std. Homo_std has 8,530 homodimers in total that has <= 30% pairwise sequence identity. It is split into a training dataset (5,975 dimers), a validation dataset (853 dimers), and a test dataset (1,702 dimers) according to the ratio of 7:1:2 to train, validate, and test DRCon.

In addition, two independent datasets (the CASP14-CAPRI dataset and DeepHomo dataset) are used to test DRCon. The CASP14-CAPRI dataset contains 7 homodimers collected from five homodimers (T1032, T1054, T1083, T1078, T1087) and two homotrimers (T1052 and T1070) that were used in the 2020 CASP14-CAPRI experiment and whose experimental structures are publicly available. From each homotrimer, we select only one homodimer with most interchain contacts.

The DeepHomo dataset used here contains 218 homodimers out of the 300 homodimers in its original version (Yan & Huang, 2021). 82 homodimers in the original DeepHomo dataset that are present in the Homo_std training dataset are removed to avoid the evaluation bias.

The statistics of the number of the dimers, the length of the dimers (i.e., the length of the monomer sequence in a homodimer), and the contact density of the dimers (i.e., the number of true interchain contacts divided by the length of the monomer in a homodimer) of the three test datasets above is reported in **Table 1**.

**Table 1.**
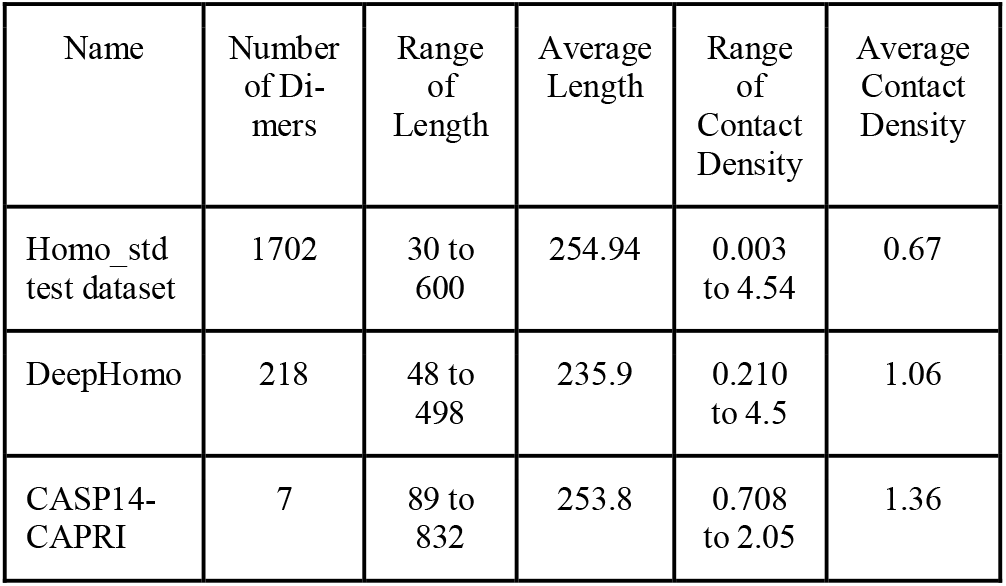
The statistics of the Homo_std test dataset, DeepHomo test dataset and CASP14-CAPRI test dataset

### 2.2 Input features

The input features for DRCon are stored in L x L x d tensors (L: length of the sequence of the monomer in a homodimer; d is the number of features for each pair of interchain residues) that describe the features of all pairs of interchain residues. Since the two chains in a homodimer are identical and interchain residue-residue coevolution features are also preserved in the multiple sequence alignment (MSA) of one chain (monomer), only the sequence of a monomer is used to generate the input features for interchain contact prediction in this work.

The number of features (d) for each interchain residue pair is 592. 49 features are the same kind of features used by DNCON2 (Adhikari et al., 2018) for intrachain contact prediction, including solvent accessibility of residues as well as interchain residue-residue coevolution features calculated from MSAs of a monomer by CCMpred (Seemayer et al., 2014) and PSICOV(Jones et al., 2012). 526 features generated from MSAs by trRosetta (Yang et al., 2020) are also used. The 8-state secondary structure prediction for each residue (i.e., 16 features for a pair of residues) made by SCRATCH (Cheng et al., 2005) is also included. Finally, a binary feature indicating if two residues form an intrachain contact (i.e., Cb-Cb atom distance is less than or equal to 8Å (Adhikari et al., 2015; Wu et al., 2021) is also used as input, which is useful for the neural network to distinguish interchain contacts from intrachain contacts. In the training phase, the intrachain contacts are derived from the true tertiary structures of monomers in the dimers. In the test phase, the intrachain contacts may be either derived from true tertiary structures of monomers in the bound state or predicted from sequences/tertiary structure models of monomers in the unbound state, depending on the experimental setting. Specifically, for the training and validation datasets, the intrachain contacts are derived from the known tertiary structures of the monomers in the homodimers (the bound state). For the test datasets, either true intrachain contacts or predicted intrachain contacts made by trRosetta or extracted from AlphaFold2 tertiary structure models in the unbound state are used to generate intrachain contact features.

Most of the 592 features above are generated from the MSAs of the monomers in the homodimers. The DNCON2’s MSA generation procedure is used to generate the MSAs for all the datasets by using HHBlits (Remmert et al., 2012) to search UniRef30_2020_02 database (Suzek et al., 2015) and Jackhmmer (Johnson et al., 2010) to search Uniref90. In addition, DeepMSA (Zhang et al., 2020) is used to generate MSAs for the CASP14-CAPRI dataset. The MSAs with more sequences generated by DNCON2 or DeepMSA are selected for the proteins in this dataset.

### 2.3 Deep learning architecture for interchain contact prediction

**Figure 1** illustrates the deep learning architecture for interchain contact prediction. The input tensor (L x L x 592) is first transformed by a block consisting of a convolutional layer and instance normalization. The instance normalization instead of the batch normalization is used because the former is better at dealing with a small batch size (Lian & Liu, 2019). The transformed tensor is then processed by 36 residual blocks containing regular convolutional layers, instance normalization, dilated convolutional layers, and residual connections. The residual connection makes the learning of deep networks more efficient and effective. The dilated convolution can capture a larger input area than the regular convolution with the same number of parameters, which has been shown to improve intrachain residue-residue distance prediction in AlphaFold1 (Senior et al., 2019).

**Figure 1.**
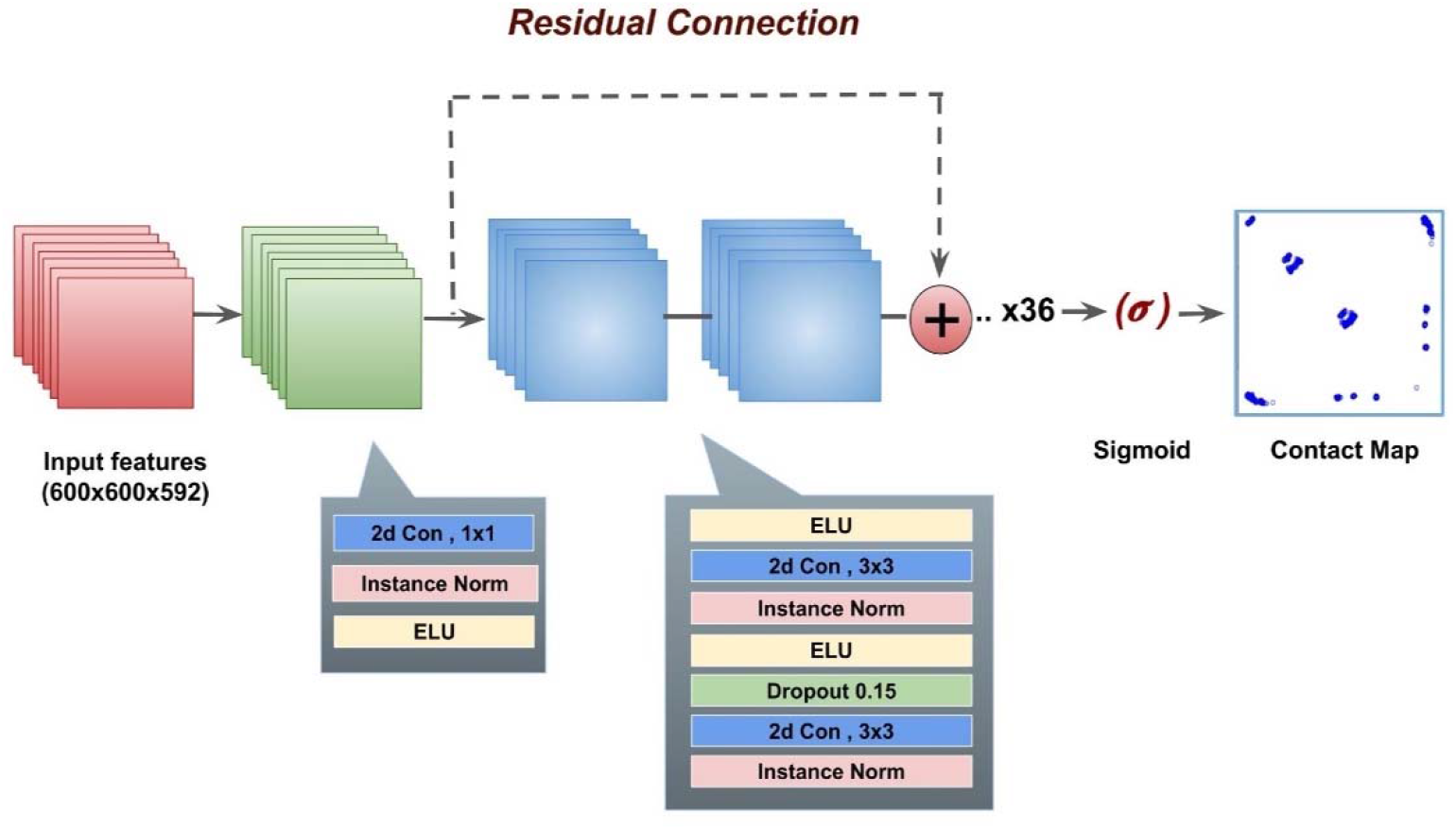
The deep learning architecture of DRCon for interchain contact prediction in homodimers. For a homodimer in which the length of the monomer sequence is L, the input is a L x L x 592 tensor. The number of input features for each pair of residues is 592. For convenience, L is set to a fixed number - 600. 0 padding is applied if L is less than 600. It is worth noting that in the prediction phase, no zero padding is used in generating the input tensor if L is greater than 600. The input is transformed to a 600 × 600 × 48 tensor using a 2D-convolutional layer which has a kernel size of 1 and uses Exponential Linear Unit (elu). The output of the convolution layer is passed through 36 residual blocks with kernel size of 3×3. Each residual block uses a 2D-convolution layer with a kernel size of 3, instance normalization and dropout of 15% probability of a neuron being ignored, followed by a dilated convolution layer without dropout. The step of the dilation in the dilated convolution layers in these blocks changes from 1, 2, 4, 8, 16 periodically. The sigmoid activation function is applied to the output of the last residual block to calculate the contact probability of each interchain residue-residue pair. The probabilities for residue pair (i,j) and residue pair (j, i) are averaged to a symmetric final contact map.

The network is trained on the Homo_std training dataset with 0.0001 learning rate and optimized with Adam (Kingma & Ba, 2017) optimizer using a batch size of 2 and the binary cross entropy as loss function. Each epoch of training the network on six 32GB NVIDIA V100 GPUs takes around 2 hours. The deep network is implemented on Pytorch and horovod (Sergeev & Del Balso, 2018) to leverage the distributed deep learning training. The deep learning model with the highest precision for top L/5 interchain contact predictions on the Homo_std validation dataset is selected as the final model for testing.

## 3 Results and Discussions

DRCon has been extensively benchmarked on three datasets: Homo_std test dataset, DeepHomo test dataset and CASP14-CAPRI dataset. The contact-level precision and the target-level accuracy rate at the various thresholds (i.e., Top 10, top L/10, top L/5, top L interchain contact predictions) are used to compare DRCon with existing methods, where L is the length of the monomer sequence in a homodimer. The contact-level precision is the number of correctly predicted contacts divided by the total number of contact predictions. And the target-level accuracy rate (Zhao & Gong, 2019) is defined as the percentage of dimers (targets) with nonzero correct interchain contact prediction when a certain number of predicted interchain contacts are evaluated.

### 3.1 Evaluation on Homo_std test dataset

We compare DRCon with DNCON2_Inter on the Homo_std test dataset. DRCon is run in the two settings. In one setting, the true intrachain contacts extracted from known tertiary structures of a monomer in each homodimer are used as input. In another setting, the intrachain contacts predicted by trRosetta are used as input. Predicted intrachain contacts are converted from the distance probabilities predicted by trRosetta. A cutoff probability of 0.5 is applied to make the conversion. The precision of top L and top 2L intrachain contact predictions made by trRosetta is 86% and 78%, respectively, indicating the quality of the intrachain contact prediction is good.

The precision of the interchain contact prediction on the Homo_std test dataset is reported in **Table 2**. The precision of DRCon in the two settings is more than twice that of DNCON2_Inter. For instance, the precision of DRCon with predicted intrachain contact prediction as input for top L/10 interchain contact prediction is 37.25%, higher than 17.32% of DNCON2_Inter. The difference is largely because DRCon is specially designed and trained to predict interchain contacts, but DNCON2_Inter is adapted from a deep learning method designed and trained to predict intrachain contacts.

**Table 2.**
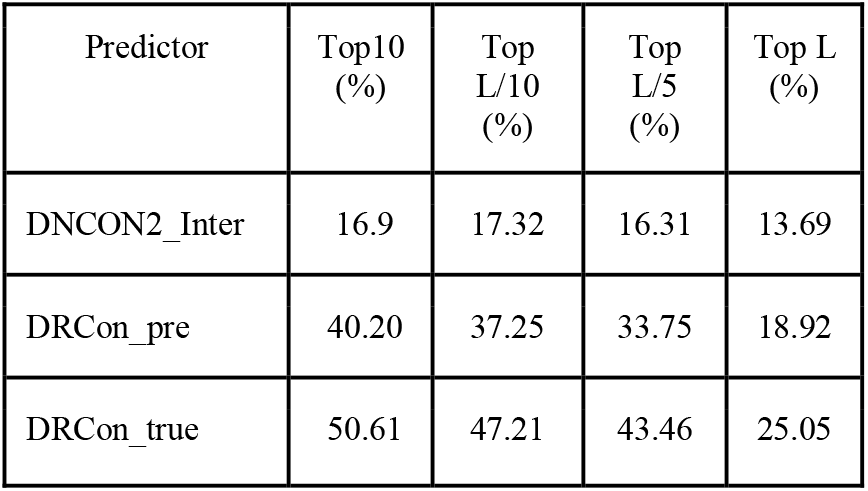
The interchain contact prediction precision of DNCON2_Inter, the DRCon with true intrachain contacts as input (DRCon_true) and DRCon with predicted intrachain contacts as input (DRCon_pre) on Homo_std test set. The precision of DNCON2_Inter is reported with its best parameter setting (relax_removal=2). L: length of a monomer in a dimer.

The precision of DRCon with predicted intrachain contacts (DRCon_pre) as input is worse than that of DRCon with true contacts by about 6 to 11 percentage points for Top 10, Top L/10, Top L/5 and Top L interchain contact predictions, indicating that more precise intrachain contact prediction (or tertiary structure prediction) of monomer leads to the higher accuracy of the interchain contact prediction. Because the predicted intrachain contacts represent the tertiary structures of monomers in the unbound state (i.e., in the free state without a binding partner) while the true intrachain contacts represent the tertiary structures in the bound state (i.e., in the state of binding with a partner in complex), the reasonable performance of DRCon_pre shows that DRCon trained on the dimers and the true tertiary structures of monomers in the bound state can work well on the predicted input intrachain contacts (or predicted tertiary structures) in the unbound state. The similar trend is also observed in the target-level prediction accuracy rate on the dataset (**Table 3**).

**Table 3.**
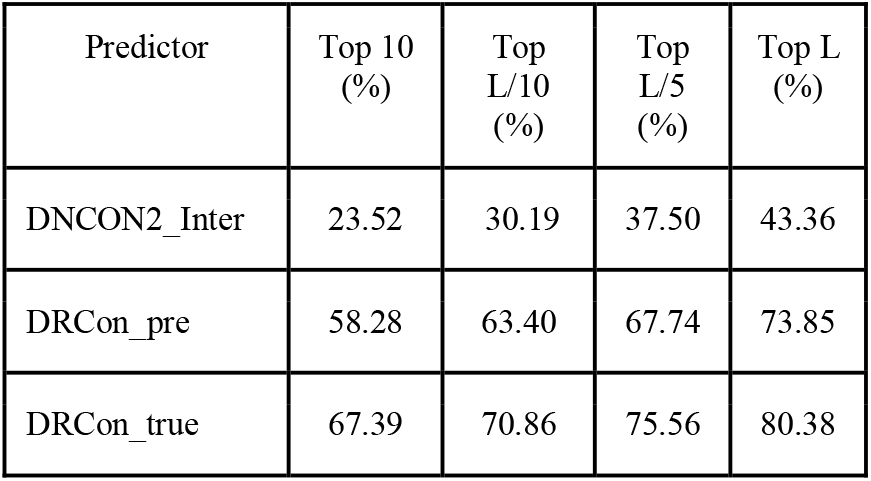
Target-level accuracy rate of DNCON2_Inter and DRCon on the Homo_std test dataset.

| Predictor | Top 10 (%) | Top L/10 (%) | Top L/5 (%) | Top L (%) |
| --- | --- | --- | --- | --- |
| DNCON2_Inter | 23.52 | 30.19 | 37.50 | 43.36 |
| DRCon_pre | 58.28 | 63.40 | 67.74 | 73.85 |
| DRCon_true | 67.39 | 70.86 | 75.56 | 80.38 |

### 3.2 Evaluation on DeepHomo test dataset

We compare DRCon and DeepHomo on the DeepHomo test dataset (see the contact-level precision and target-level accuracy rate in **Table 4** and **Table 5** respectively). For a fair comparison, we use the same tertiary structures of monomers in the homodimers provided by the DeepHomo server to extract the intrachain contacts as input for DRCon and for the DeepHomo server itself to make interchain contact predictions. The interchain contacts predicted by DeepHomo consist of only the upper triangle of the interchain contact map. They are converted to a diagonally symmetric full contact map for evaluation as DeepHomo assumes the contact map is of C2-symmetry. DRCon performs better than DeepHomo in terms of contact-level precision and target-level accuracy rate at all the thresholds except for the target-level accuracy rate of top L contact predictions. For instance, the contact-level precision and target-level accuracy of DRCon for top L/10 interchain contact prediction is 50.17% and 76.15%, higher than 38.74% and 70.77% of DeepHomo.

**Table 4.**
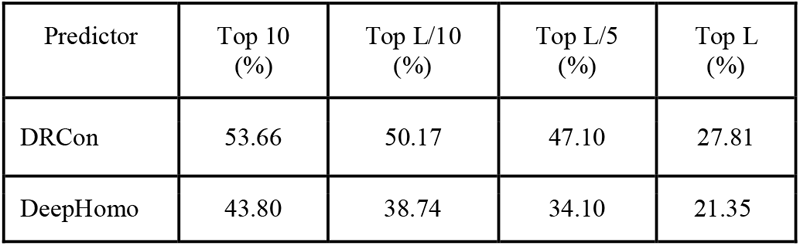
The interchain contact prediction precision of DRCon and DeepHomo on the DeepHomo test dataset

| Predictor | Top 10 (%) | Top L/10 (%) | Top L/5 (%) | Top L (%) |
| --- | --- | --- | --- | --- |
| DRCon | 53.66 | 50.17 | 47.10 | 27.81 |
| DeepHomo | 43.80 | 38.74 | 34.10 | 21.35 |

**Table 5.** The target-level accuracy rate of DRCon and DeepHomo on the DeepHomo test dataset

| Predictor | Top 10 (%) | Top L/10 (%) | Top L/5 (%) | Top L (%) |
| --- | --- | --- | --- | --- |
| DRCon | 72.95 | 76.15 | 80.73 | 86.69 |
| DeepHomo | 66.67 | 70.77 | 77.17 | 87.61 |

### 3.3 Evaluation on CASP14-CAPRI dataset using true or predicted tertiary structures as input

We compare DRCon and DeepHomo on the CASP14-CAPRI dataset. Only the contact-level precision is used to evaluate them because only 7 targets are not sufficient to reliably estimate the target-level accuracy rate. DRCon is run in the two settings (the ideal setting and the realistic setting). In the ideal setting (DRCon_true), the known tertiary structures of the monomers in the homodimers are used to generate the true intrachain contacts as input for DRCon. In the realistic setting (DRCon_alpha), the tertiary structures of the monomers predicted by AlphaFold2 (Jumper et al., 2021) are used to generate the interchain contacts for DRCon. The AlphaFold2 model with the highest confidence is used for each target. The TM-scores (Y. Zhang & Skolnick, 2004) of the tertiary structures for the 7 targets (T1032, T1052, T1054, T1070, T1078, T1083, T1087) predicted by AlphaFold2 is 0.708, 0.672, 0.924, 0.491, 0.985, 0.870, and 0.977. The average TM-score is 0.840.

The precision of interchain contact predictions of DRCon and DeepHomo are shown in **Table 6**. The precision of both DRCon_true and DRCon_alpha is substantially higher than that of DeepHomo at all the thresholds. For instance, for top L/10 interchain contact predictions, the precision of DRCon_true and DRCon_alpha is 30.6% and 26.57%, more than double 11.63% of DeepHomo. DRCon_true performs better than DRCon_alpha, indicating that more accurate intrachain contact input leads to better interchain contact prediction.

**Table 6.** The precision of DRCon and DeepHomo on the CASP14-CAPRI test dataset. DRCon_true and DeepHomo use the true tertiary structures of monomers in the bound state to extract intrachain contacts as input. DRCon_alpha uses the tertiary structures predicted by AlphaFold2 in the unbound state to extract intrachain contacts as input.

| Predictor | Top 10 (%) | Top L/10 (%) | Top L/5 (%) |
| --- | --- | --- | --- |
| DRCon_true | 32.85 | 30.06 | 24.81 |
| DRCon_alpha | 27.14 | 26.57 | 20.76 |
| DeepHomo | 11.42 | 11.63 | 13.09 |

The precision of top L/5 contact predictions for each target made by the methods is illustrated in **Figure 2**. T1032 and T1052 are classified by CAPRI as easy targets that have dimer structure templates in PDB, while T1054, T1070, T1078, T1083, and T1087 are classified as difficult targets that do not have good dimer structure templates. DRCon_true, DRCon_alpha and DeepHomo perform better on T1032, T1078, and T1087 than on the other four targets. Both DRCon_true and DRCon_alpha outperform DeepHomo on all but one target (T1078). However, all the methods perform poorly on two dimers T1052 and T1070 extracted from homotrimers, indicating it may be more challenging for the methods trained only on homodimers to predict dimerization interactions in homo multimers consisting of more than two protein chains. All the methods failed to predict any interchain contact for T1083. DRCon_true performs better than or similarly to DRCon_alpha on all but one target (T1087), confirming better intrachain contact input (or tertiary structure input) generally leads to better interchain contact prediction. The tertiary structure predicted by AlphaFold for T1087 is very similar to the true structure (TM-score of the AlphaFold structure = 0.977), suggesting that the difference in their performance on this target may be mostly due to the random variation in the input instead of the quality of intrachain contact input.

**Figure 2.**
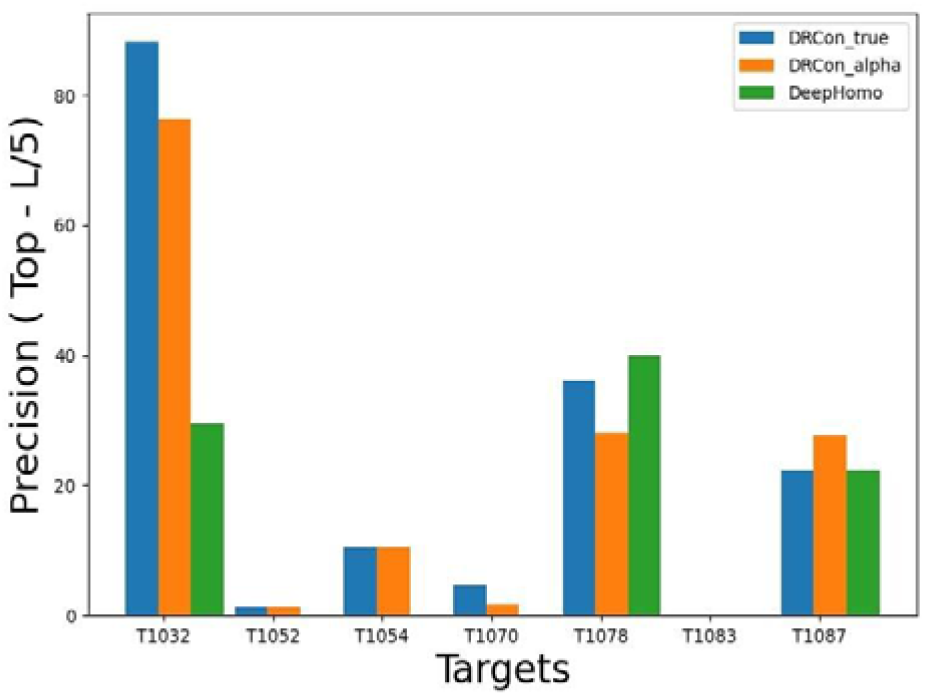
The precision of top L/5 interchain contact predictions for each of 7 targets.

### 3.4 Effect of contact density on interchain contact prediction

We investigate the interchain contact density in a dimer with the precision of the interchain contact prediction on the Homo_std test dataset. The Pearson’s correlation coefficient between the precision of top L/5 interchain contacts and contact density is 0.4211, indicating a moderate correlation between the two. The lowest average precision (a little over 4%) is recorded for targets with the low contact density between 0 and 0.25, indicating that when the interchain contact map is very sparse, the prediction is generally difficult.

### 3.5 A case study of applying interchain contact prediction to build quaternary structure

**Figure 3** visualizes the top L/5 interchain contact predictions for a target (PDB code: 1DR0) from the Homo_std test dataset and the quaternary structure reconstructed from the interchain contacts predicted by DRcon and the known tertiary structure of a chain in the dimer. The quaternary structure is built by GD (Soltanikazemi et al., 2021), which applies the gradient descent optimization to build quaternary structures by using interchain contacts as distance restraints.

**Figure 3.**
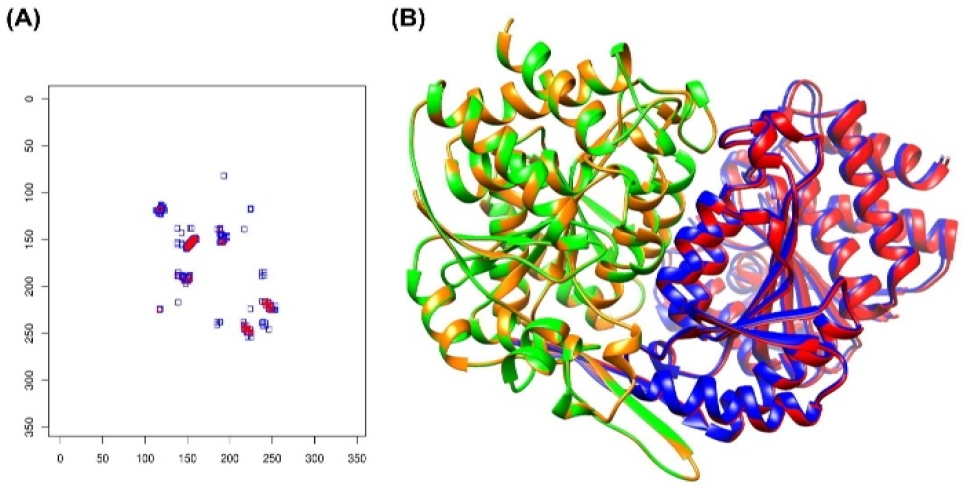
**(A)** The predicted and true contact maps of target 1DR0. The top L/5 predicted contacts (red dots) and true contacts (blue dots) are plotted. Most predicted contacts overlap with the true contacts, indicating a high contact prediction precision. **(B)** The superimposition of the true quaternary structure (chain A in red and chain B in green) and the predicted quaternary structure (chain A in blue and chain B in orange). The two quaternary structures are quite similar.

It is shown in **Figure 3A** that most of the interchain contact predictions overlap with the true interchain contacts, indicating a high prediction precision. Indeed, the precision of top L/5 and top L contact predictions is 100% and 75%, respectively. The quaternary structure reconstructed from the predicted interchain contacts is also very similar to the native structure (**Figure 3B**). The TM-score of the predicted quaternary structure in comparison with the true quaternary structure is 0.99. TMalign (Y. Zhang & Skolnick, 2005) is used to calculate the TM-score. The predicted quaternary structure has a fraction of the native contacts (F_nat_) of 0.88, interface RMSD (iRMS: root mean square displacement of inter-protein heavy atoms that are within 10 Å) of 0.3 Å, ligand RMS (LRMS) of 0.83 Å and a DockQ score of 0.95. F_nat_, iRMS, LRMS, and DockQ score of the predicted quaternary structure are calculated against the true quaternary structure by DockQ (Basu & Wallner, 2016). A DockQ score of 0.8 indicates a high-quality quaternary structure prediction.

## Conclusion and Future Work

In this work, we develop a deep network (DRCon) consisting of residual connections, regular and dilated convolutions and instance normalizations to predict interchain homodimers from sequence and structural features of monomers in homodimers. DRCon trained on known homodimer structures can predict interchain contacts well. Moreover, DRCon is robust against the errors in input tertiary structures or intrachain contacts of monomers. It maintains the reasonable prediction precision when predicted tertiary structures of monomers in the unbound state instead of true tertiary structures in the bound state are used as input. The work demonstrates that deep learning methods specially designed for interchain contact prediction can be trained on known homodimer structures to substantially improve the prediction of interchain residue-residue contacts as what had happened in protein tertiary structure prediction. In the future, we plan to further improve the deep learning architecture, input features, and training strategies to improve interchain contact prediction. We will generalize the method to predict interchain residue-residue distances. Moreover, we plan to develop similar methods to predict interchain contacts and distances in heterodimers and generalize them to multimers consisting of more than two chains.

## Acknowledgements

The work was partly supported by the Department of Energy, USA (DE-AR0001213, DE-SC0020400 and DE-SC0021303), National Science Foundation (DBI1759934 and IIS1763246), National Institutes of Health (R01GM093123), and the Thompson Missouri Distinguished Professorship. The project leveraged the Oak Ridge Leadership Computing Facility, which is a DOE Office of Science User Facility supported under Contract DE-AC05-00OR22725.

